# Protective efficacy of COVAXIN® against Delta and Omicron variants in hamster model

**DOI:** 10.1101/2022.06.14.496021

**Authors:** Pragya D Yadav, Sreelekshmy Mohandas, Anita Shete, Gajanan Sapkal, Gururaj Deshpande, Abhimanyu Kumar, Kundan Wakchaure, Hitesh Dighe, Rajlaxmi Jain, Brunda Ganneru, Jyoti Yemul, Pranita Gawande, Krishna Mohan, Priya Abraham

## Abstract

The immunity acquired after natural infection or vaccinations against SARS-CoV-2 tend to wane with time. Vaccine effectiveness also varies with the variant of infection. Here, we compared the protective efficacy of COVAXIN® following 2 and 3 dose immunizations against the Delta variant and also studied the efficacy of COVAXIN® against Omicron variants in a Syrian hamster model. The antibody response, clinical observations, viral load reduction and lung disease severity after virus challenge were studied. Protective response in terms of the reduction in lung viral load and lung lesions were observed in both the 2 dose as well as 3 doses COVAXIN® immunized group when compared to placebo group following the Delta variant challenge. In spite of the comparable neutralizing antibody response against the homologous vaccine strain in both the 2 dose and 3 dose immunized groups, considerable reduction in the lung disease severity was observed in the 3 dose immunized group post Delta variant challenge indicating the involvement of cell mediated immune response also in protection. In the vaccine efficacy study against the Omicron variants i.e., BA.1 and BA.2, lesser virus shedding, lung viral load and lung disease severity were observed in the immunized groups in comparison to the placebo groups. The present study shows that administration of COVAXIN® booster dose will enhance the vaccine effectiveness against the Delta variant infection and give protection against the Omicron variants BA.1.1 and BA.2.

## Introduction

SARS-CoV-2 has evolved to numerous variants in the last two years posing challenge to the mankind. The immunity generated by the natural infection or vaccination tends to wane with the time.^1,2^ Waning of immune responses post primary 2 dose series of vaccination has been reported for the available COVID-19 vaccines.^3^ The newly evolving variants have also posed challenge to the acquired immunity with their immune escape properties. Thus the Variants of Concerns (VoCs) which have evolved till date managed to spread globally even after good vaccination coverage. Vaccine break through infections are being reported worldwide.^4^ This necessitates the importance of constant monitoring of the virus genomic changes and the properties of the evolving variants. The decline of immunity varies with vaccine products as well as the target vaccination group. World Health Organization has recommended for the administration of additional dose for the inactivated COVID-19 vaccines like CoronaVac and BBIP.^5,6^ Booster doses have shown to improve the efficacies of many COVID-19 vaccines in clinical trials and has been authorized by regulatory authorities in many countries.^7–10^

Vaccine effectiveness differed for disease severity, symptomatic disease and infection for the different variant of Concerns.^11^ In case of the Alpha variant, the protection was retained against the above mentioned outcomes by the vaccines. With the Beta and Delta variant, the vaccines showed reduced protection against symptomatic disease whereas disease severity was found reduced in the vaccinated population. Of the 5 designated Variant of Concerns, Omicron variant is in current circulation throughout the world and the other variants constitutes less than 1% in prevalence. The Omicron lineage has been further divided into multiple sub lineages of which, few like BA.1, BA.2, BA.4 and BA.5 have shown growth advantage. The Omicron variant is known to escape many therapeutic/vaccine elicited neutralizing antibodies and have shown reduced effectiveness following the 2 dose regimen of many COVID-19 vaccines.^12,13^

COVAXIN® is an inactivated SARS-CoV-2 whole virion vaccine licensed in India and 13 other countries.^4^ The vaccine has shown immunogenicity and protective efficacy against SARS-CoV-2 B.1 variant possessing the D614G mutation in laboratory animal studies.^14,15^ After the successful completion of the Phase I to III human clinical trials, vaccine has received the WHO emergency authorization approval.^16^ The vaccine efficacy reported was 77.8% of symptomatic COVID-19.^16^ With the emergence of SARS-CoV-2 variants, multiple *in vitro* studies were conducted to understand the neutralization potential of the COVAXIN® induced immune responses.^17,18^ Neutralization potential of the COVAXIN® vaccinee sera has been demonstrated with the Alpha, Kappa, Beta, Delta and Omicron variants. But the neutralizing antibody titres against the VoCs were found reduced in comparison to the ancestral B.1 variant.^17–19^ Among these variants, Omicron was least effectively neutralized in the *in vitro* assays.^20^ As the cellular immune response is also a major part of the protective immune response mechanism of the body, the mere measurement of neutralizing antibodies in the sera will not tell us the whole story. The vaccine efficacy data of COVAXIN® against Omicron variant is not available.

As the vaccine responses tend to vary with the variant divergence, and the immune memory, the approved vaccines should be monitored for their protective efficacy against newly evolving variants. Here, we have performed an animal model challenge study to assess the protective efficacy of COVAXIN® against the VoCs *ie*., Delta and Omicron variants. Syrian hamster model is a widely used animal model for SARS-CoV-2 preclinical research worldwide.^21,22^ In the first study, we have compared the protective efficacy in 2 and 3 dose vaccinated in hamsters with the Delta variant challenge and in the second study, protective response was assessed the against Omicron variants following 3 dose vaccinations.

## Results

### Antibody response in hamsters post vaccination and Delta challenge

In the present study, we compared the protective efficacy of 2 and 3 doses of COVAXIN® against the infection of SARS-CoV-2 Delta variant in hamsters (Figure 1a). In all the immunized animals, 14 days post second or third dose, seroconversion was observed (Figure 1b). Anti-RBD IgG levels [mean Optical Density (OD) ± Standard Deviation (SD) =1.27 ± 0.60 in 3 dose group, 0.77 ± 0.42 in 2 dose group, serum dilution: 1: 100] and geometric mean PRNT titres (GMT) (GM ± SD = 604.5 ± 2.4 in 2 dose group and 944.5 ± 4.05 in 3 dose group) were higher in the 3 dose immunized group (Figure 1c, 1d).

**Figure 1:**
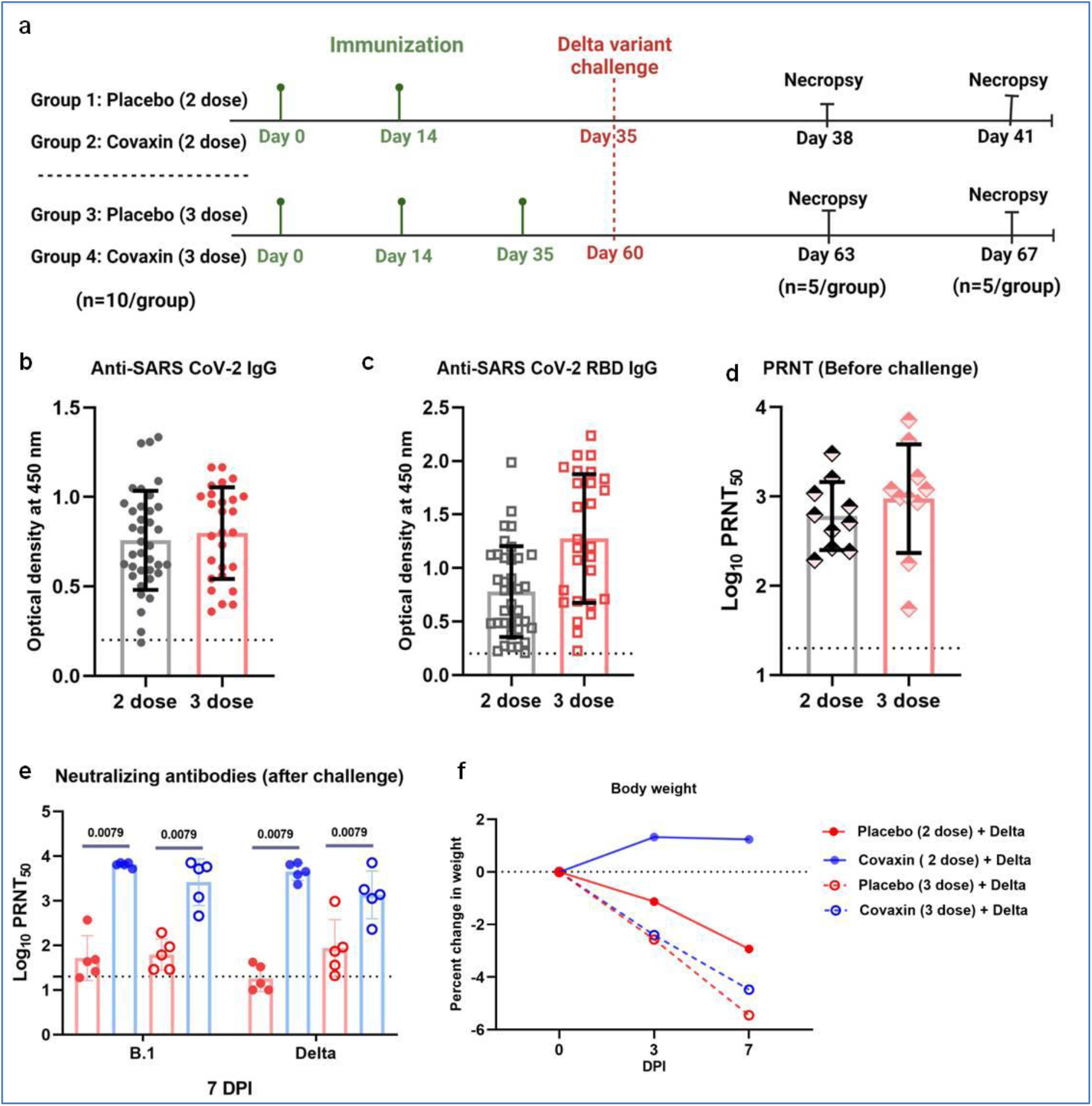
Study design, antibody response and body weight changes in hamsters following COVAXIN® immunization and Delta variant virus challenge. a) Summary of the study design of protective efficacy of COVAXIN® against Delta variant infection. Antibody response in COVAXIN® immunized hamsters two weeks post the last dose of vaccination as measured by a) Anti-SARS-CoV-2 IgG ELISA and b) Anti-SARS-CoV-2 RBD IgG ELISA. Mean along with standard deviation ids plotted on the graph, n=20/ group). The dotted lines indicate the limit of detection of the assay. c) Neutralizing antibody response in hamsters against the SARS-CoV-2 B.1 variant before Delta virus challenge (n=10/group) and e) after Delta variant challenge on 7 DPI (n= 5/group). The dotted lines indicate the limit of detection of the assay. Geometric mean along with the standard deviation is plotted on the graph. The significant differences between the experimental groups were assessed using Mann Whitney test and the p value is plotted above the bars of the respective groups. P values less than 0.05 were considered statistically significant. f) The mean body weight change in hamsters after Delta variant challenge on 3 (n= 10/ group) and 7 DPI (n= 5/ group).

The GMT against the vaccine strain after Delta variant challenge showed a 10 fold rise in 2 dose group (GM ± SD= 2594 ± 3.32) and a 2.7 fold rise in the 3 dose group (GM ± SD 6420 ± 1.13) when compared with the titres before challenge in the immunized group. The immunized animals showed a significant rise in the protective antibody titres in comparison to the placebo group too at 7 DPI (Figure 1e). The GMT against the BBV152 vaccine strain i.e., the B.1 variant was 123 fold and 41 fold higher in the 2 and 3 dose COVAXIN® group than the placebo group respectively. Similarly against Delta variant, GMT was 250 and 15 times higher than placebo in the 2 and 3 dose vaccinated group respectively. The neutralizing antibody levels were comparable against both the Delta and the ancestral B.1 variant.

### Protection against SARS-CoV-2 Delta variant challenge

There was no significant change in the mean body weight loss observed in the immunized groups (Figure 1f). The viral RNA shed through the throat and nasal cavity was found markedly reduced in two/three dose vaccinated animals in comparison to the placebo groups (Figure 2a-d). By 7DPI, viral RNA clearance from the throat swab and nasal wash was observed in the vaccinated groups. The viral load in the nasal wash were also lower in the vaccinated groups, although significant difference could only be seen on 3 and 7 DPI of the 2 dose group (Figure 2e).

**Figure 2:**
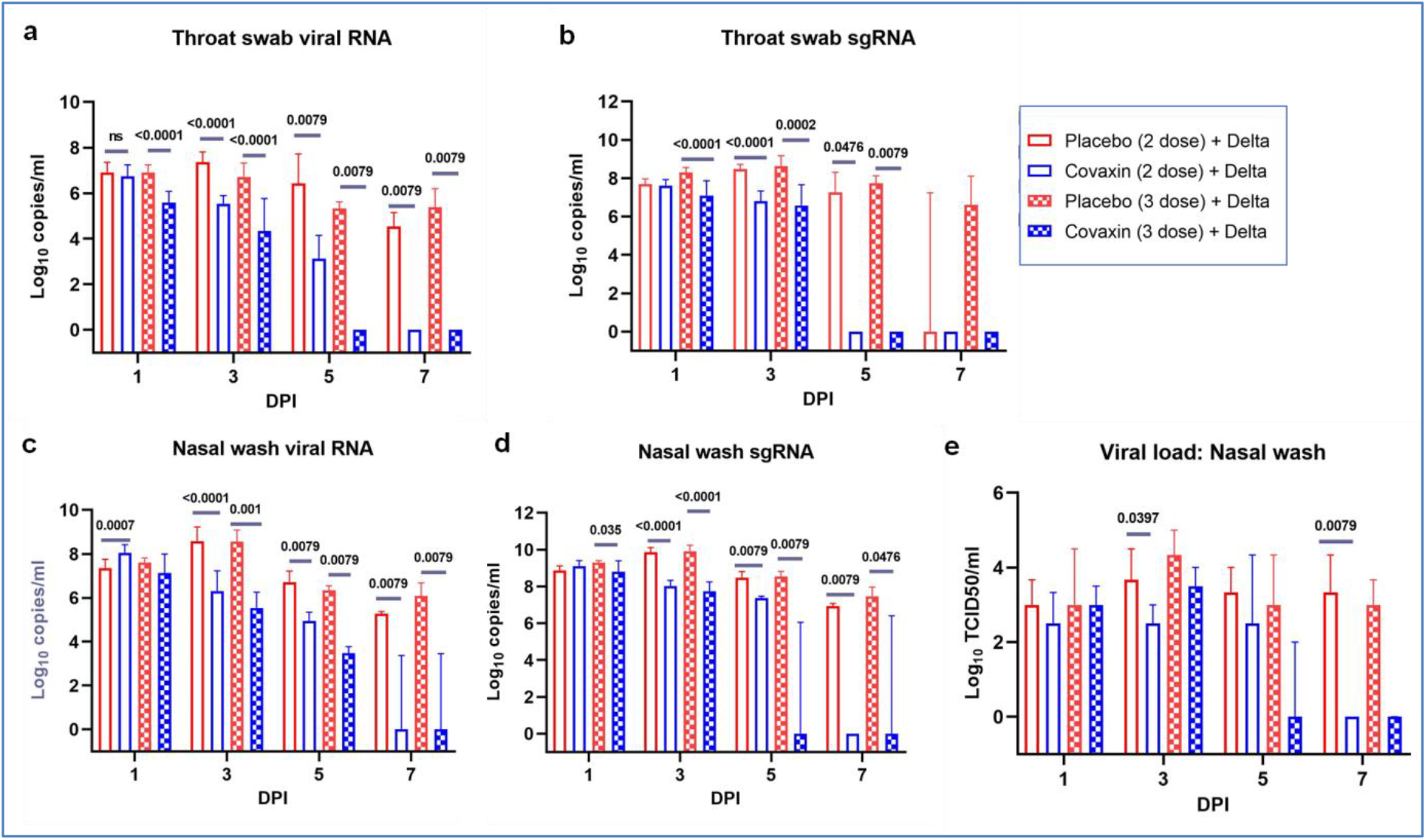
Virus shedding in COVAXIN® immunized hamsters after Delta variant challenge on 1, 3, 5 and 7 DPI. a) Viral RNA and b) subgenomic RNA levels in the throat swab samples of the immunized and placebo groups of hamsters. c) Viral RNA, d) subgenomic RNA and e) virus titres in the nasal wash samples of the immunized and placebo groups of hamsters. The median along with 95% confidence interval is plotted on the graph. The significant differences between the experimental groups (n= 10/group on 1 and 3 DPI, n= 5 /group on 5 and 7 DPI) were assessed using Mann Whitney test and the p value is plotted above the bars of the respective groups. P values less than 0.05 were considered statistically significant.

In the nasal turbinates samples, the viral RNA and subgenomic (sg) RNA were significantly reduced on 3 DPI, whereas sgRNA levels were similar on the 7DPI in both the 2 and 3 dose vaccinated groups in comparison to placebo. On virus titration, the virus titres in the 3 dose vaccinated group were found reduced on 3 and 7 DPI, whereas the 2 dose group showed similar viral titres as that of the placebo group. Lungs samples collected on 3 and 7 DPI showed significant reduction in the viral RNA and live virus titres in the vaccinated groups (Figure 3d-f).

**Figure 3:**
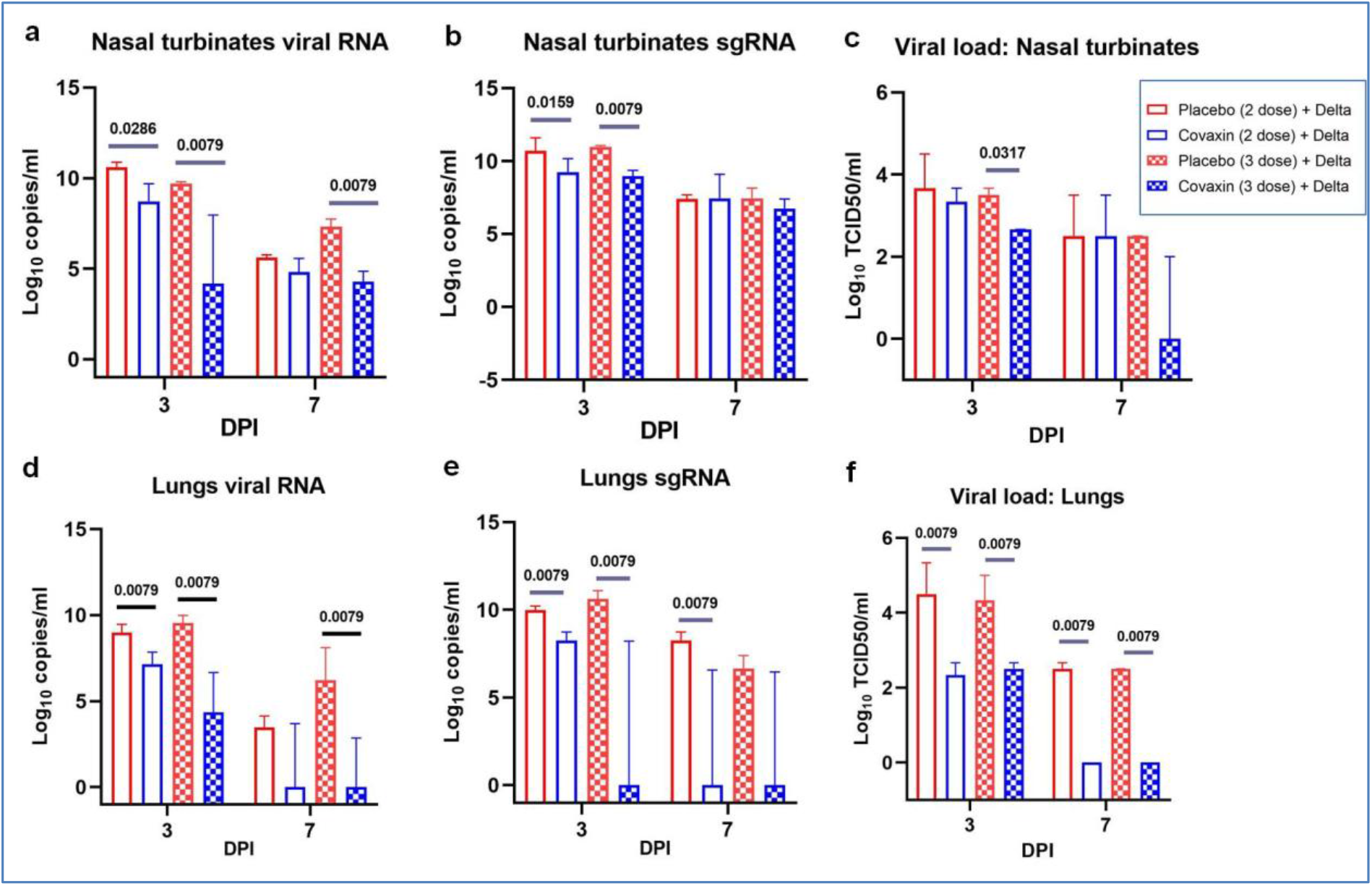
Viral load in the respiratory organs of the COVAXIN® immunized hamsters after Delta variant challenge on 3 and 7 DPI. a) Viral RNA and b) subgenomic RNA levels in the throat swab samples of the immunized and placebo groups of hamsters. c) Viral RNA, d) subgenomic RNA and e) virus titres in the nasal wash samples of the immunized and placebo groups of hamsters. The median along with 95% confidence interval is plotted on the graph. The significant differences between the experimental groups (n=5 /group) were assessed using Mann Whitney test and the p value is plotted above the bars of the respective groups. P values less than 0.05 were considered statistically significant.

**Figure 4.**
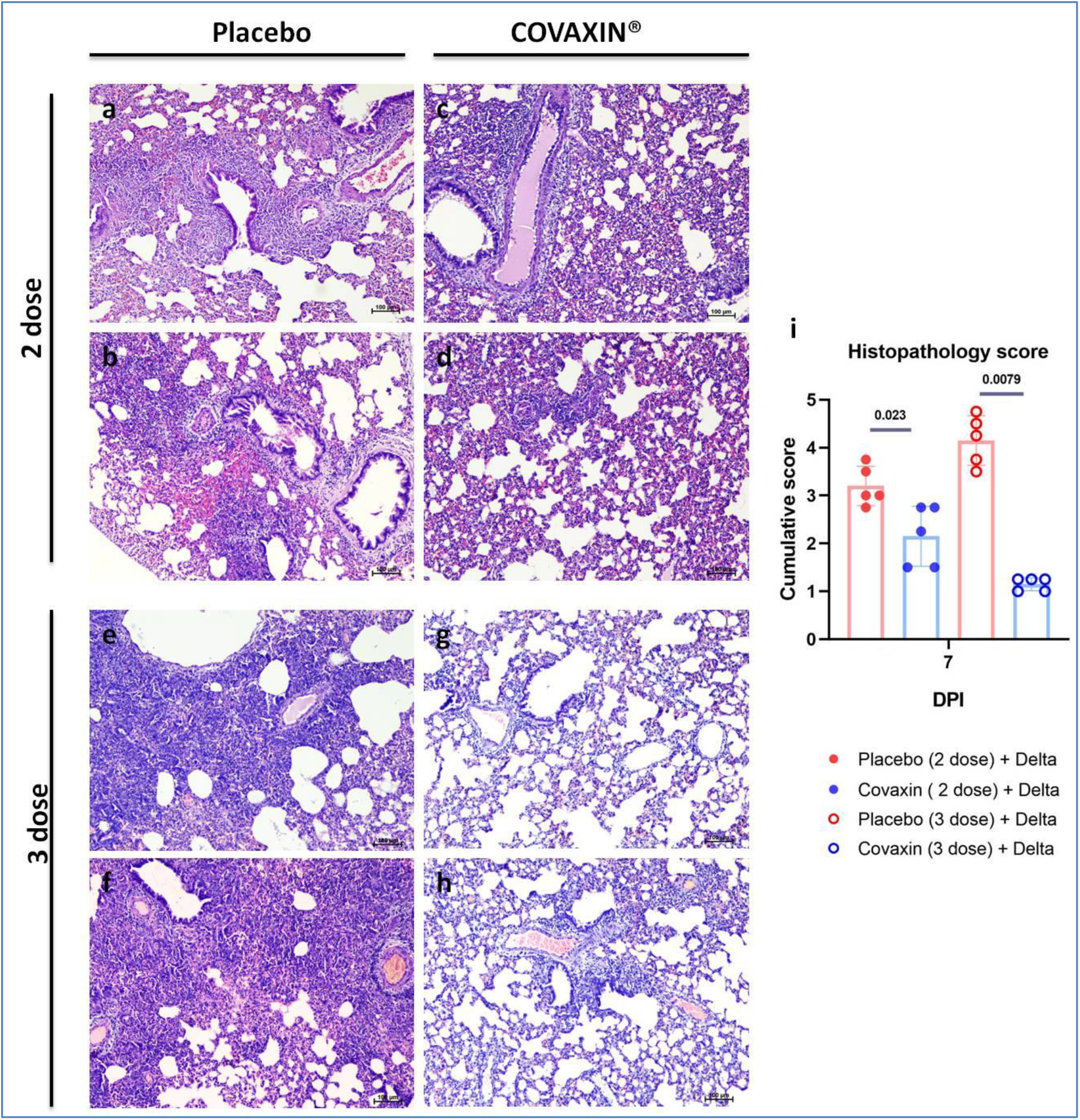
Histopathological changes in lungs of hamsters after SARS-CoV-2 Delta variant challenge. Interstitial pneumonia observed in a, b) placebo group, c,d) 2 dose immunized group and e,f) placebo (3 dose) group. The lungs of hamsters from the 3 dose COVAXIN® immunized group showing g) normal lung histology and h) mild perivasular inflammatory cell infiltration on 7 DPI. i) Cumulative histopathological score of the lung lesions post Delta variant infection in hamsters. The mean along with standard deviation is plotted on the graph. The significant differences between the experimental groups (n=5 /group) were assessed using Mann Whitney test and the p value is plotted above the bars of the respective groups. P values less than 0.05 were considered statistically significant.

The histopathological changes like perivascular and peribronchial inflammatory cell infiltration, alveolar haemorrhages, pneumocyte hyperplasia and septal thickening were observed in the Delta variant challenged animals. The lesions of lesser severity were observed in the 2 dose vaccinated groups. In the 3 dose immunized group, histopathological changes observed were minimal with focal inflammatory cell infiltration or alveolar damage compared to the placebo group.

### Protection against Omicron variant challenge

We performed a virus challenge study using the Omicron variants ie., BA.1.1 and BA.2 in hamsters following 3 dose immunization (Figure 5a). The immunized animals showed anti-SARS-CoV-2 and anti-RBD IgG response as well as high neutralizing antibody titres against B.1 (vaccine strain) variant (GMT ± SD = 2704 ± 1.618) before virus challenge (Supplementary figure 1). The body weight loss was minimal in COVAXIN® immunized hamsters infected with BA.1.1 and BA.2 (Figure 5b). The placebo group infected with BA.1.1 showed a mean weight loss of -18 ± 2.5 and weight loss was not observed in the BA.2 infected hamsters of both vaccinated and placebo group,.

**Figure 5.**
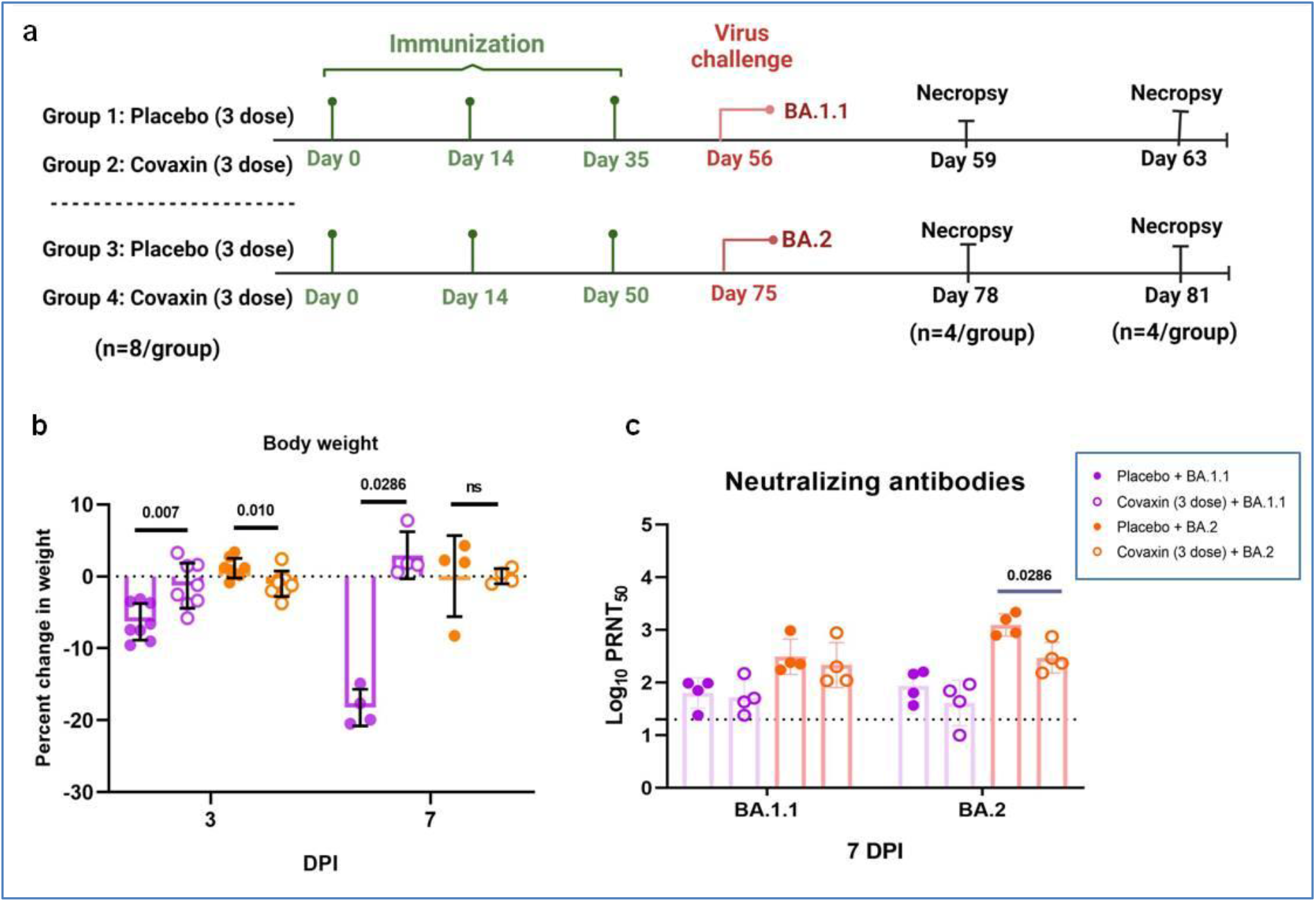
Study design, antibody response and body weight changes in hamsters following Omicron variant virus challenge. a) Summary of the study design of protective efficacy of COVAXIN® against Omicron variant infection. b) Body weight changes in hamsters after Omicron variant challenge on 3 (n= 8/ group) and 7 DPI (n= 4/ group). The mean along with standard deviation is plotted on the graph. C) Neutralizing antibody response in COVAXIN® immunized hamsters after Omicron variant challenge on 7 DPI (n= 5/group) against BA.1.1 and BA.2. The dotted lines indicate the limit of detection of the assay. The geometric mean along with standard deviation is plotted on the graph. The significant differences between the experimental groups were assessed using Mann Whitney test and the p value is plotted above the bars of the respective groups. P values less than 0.05 were considered statistically significant. ns= non significant.

The neutralizing antibody response against BA.1.1 was less in comparison to the BA.2 in both the experimental groups. The GMT was 52.92 ± 2.14 and 215.4 ± 2.67 in the BA.1.1 and BA.2 infected group against the BA.1.1 variant on 7DPI respectively. The GMT were comparable in the immunized and placebo groups on 7 DPI against BA.1.1 variant. GMT was 41.14 ± 2.69 and 296.9 ± 1.96 in the vaccinated groups against BA.2 variant after BA.1.1 and BA.2 infection (Figure 5c).

The virus shedding through nasal and oral cavity was significantly reduced in the vaccinated groups (Figure 6). Viral RNA and sgRNA levels in the nasal turbinates and lungs were also significantly reduced in the immunized hamsters after Omicron infection on day 3 and 7 after virus infection. Viral load in the nasal turbinates were comparable on 3 and 7 DPI whereas the titres in the lungs were significantly reduced on both the time points in both the vaccine groups.

**Figure 6.**
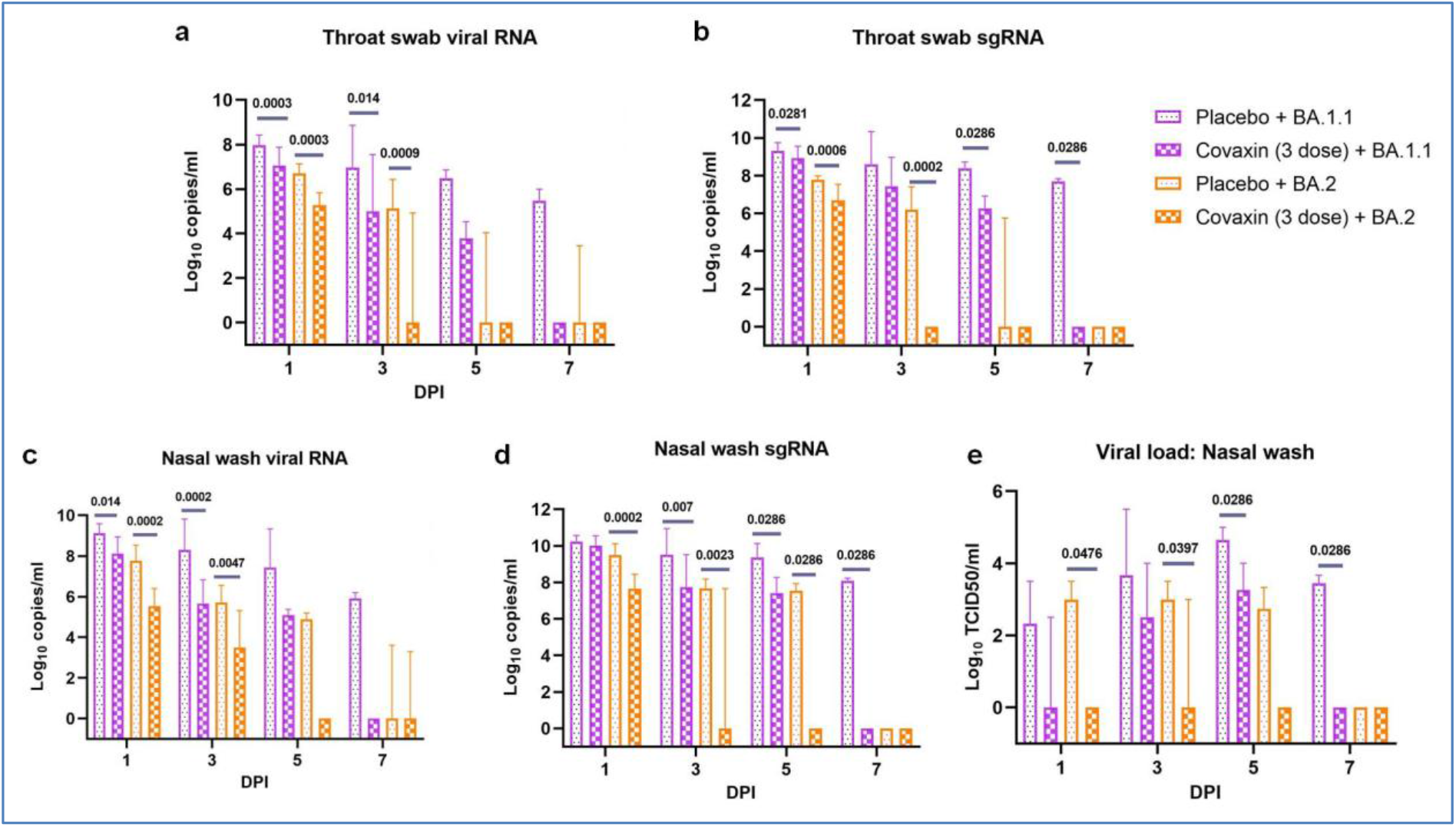
Virus shedding in COVAXIN® immunized hamsters after Omicron variant challenge on 1, 3, 5 and 7 DPI. a) Viral RNA and b) subgenomic RNA levels in the throat swab samples of the immunized and placebo groups of hamsters. c) Viral RNA, d) subgenomic RNA and e) virus titres in the nasal wash samples of the immunized and placebo groups of hamsters. The median along with 95% confidence interval is plotted on the graph. The significant differences between the experimental groups (n= 8/group on 1 and 3 DPI, n= 4 /group on 5 and 7 DPI) were assessed using Mann Whitney test and the p value is plotted above the bars of the respective groups. P values less than 0.05 were considered statistically significant.

**Figure 7.**
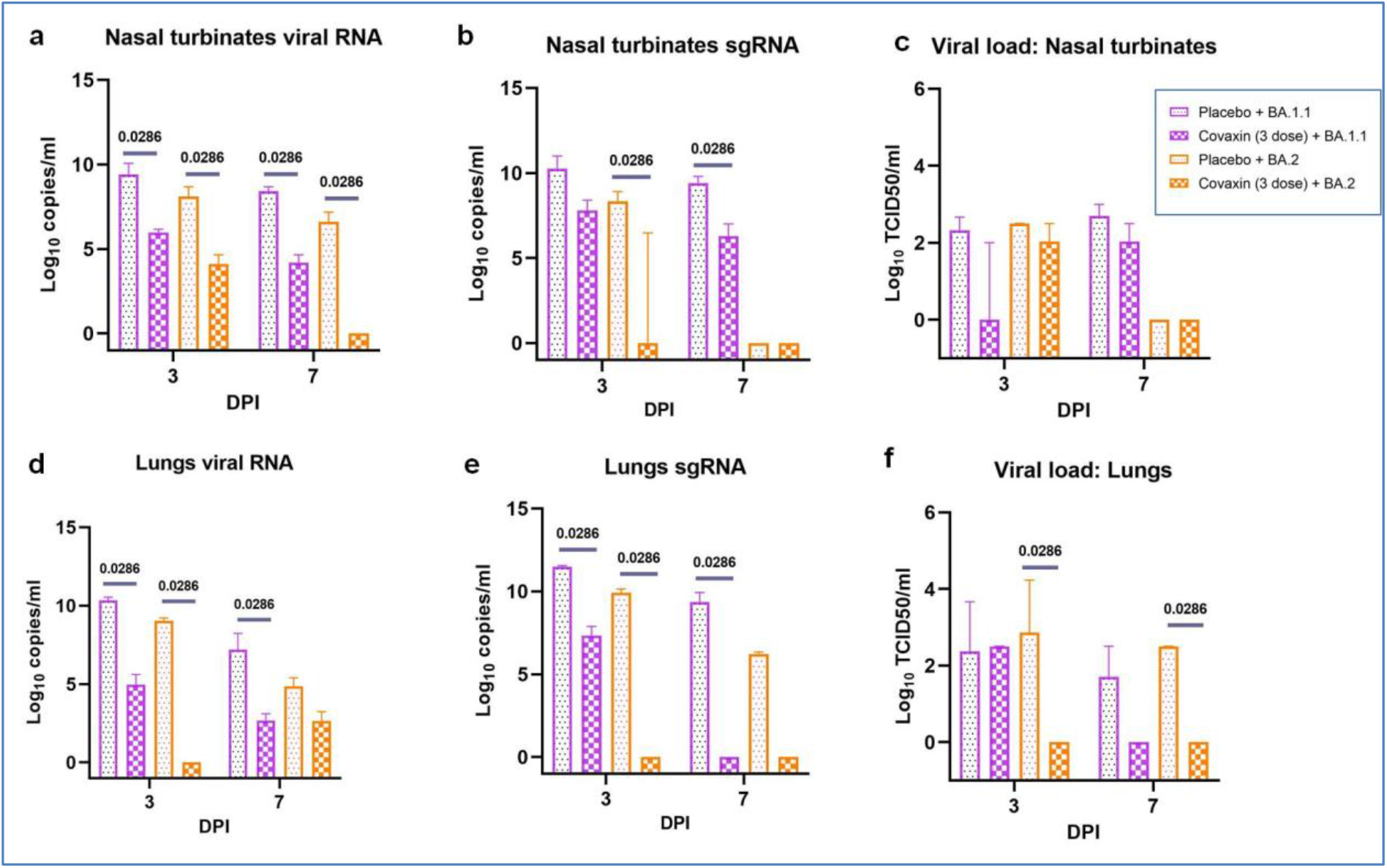
Viral load in the respiratory organs of the COVAXIN® immunized hamsters after Omicron variant challenge on 3 and 7 DPI. a) Viral RNA and b) subgenomic RNA levels in the throat swab samples of the immunized and placebo groups of hamsters. c) Viral RNA, d) subgenomic RNA and e) virus titres in the nasal wash samples of the immunized and placebo groups of hamsters. The median along with 95% confidence interval is plotted on the graph. The significant differences between the experimental groups (n=4 /group) were assessed using Mann Whitney test and the p value is plotted above the bars of the respective groups. P values less than 0.05 were considered statistically significant.

**Figure 7.**
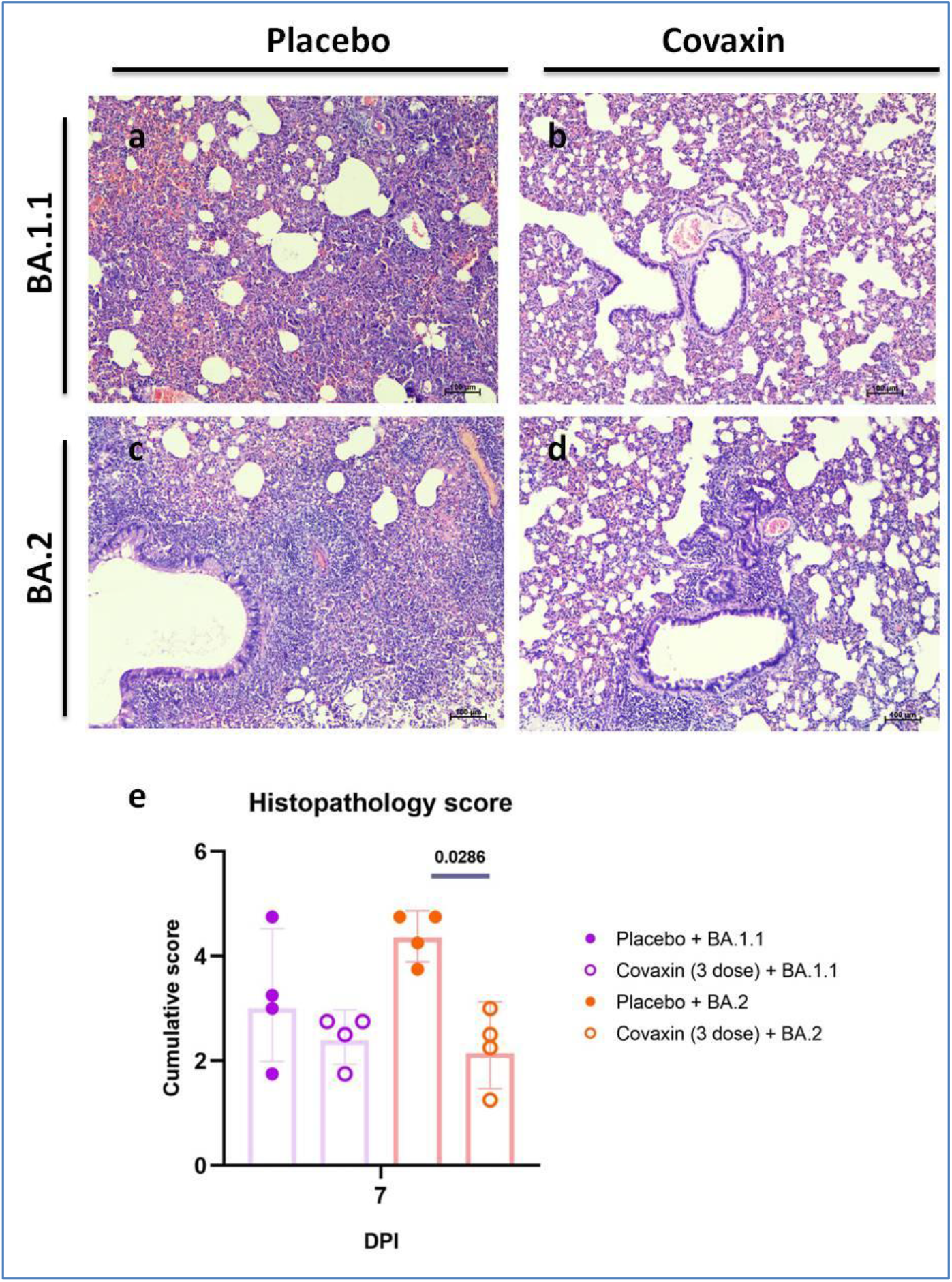
Histopathological changes in lungs of hamsters after SARS-CoV-2 Omicron variant challenge. The lungs of BA.1.1 infected hamsters of the a) placebo group showing interstitial pneumonia and b) immunized group showing alveolar capillary congestion on 7DPI. The lungs of BA.2 infected hamsters of the c) placebo group showing interstitial pneumonia characterized by diffuse alveolar septal thickening, exudates in alveoli and peribronchial inflammatory cell infiltration and d) immunized group showing alveolar capillary congestion and a foci of peribronchial inflammatory cell infiltration. i) Cumulative histopathological score of the lung lesions post Omicron variant infection in hamsters. The mean along with standard deviation is plotted on the graph. The significant differences between the experimental groups (n=4 /group) were assessed using Mann Whitney test and the p value is plotted above the bars of the respective groups. P values less than 0.05 were considered statistically significant.

Alveolar interstitial pneumonia along with peribronchial infiltration was observed in the hamsters of the placebo group infected with the Omicron variants, BA.1 and BA.2. In the vaccinated groups lesions observed were mostly focal in distribution. The lesion severity following the BA.1.1 infection was found lesser in the vaccinated hamsters with a mean (± SD) cumulative score of 2.438 (± 0.47) against a score of 3.18 (± 1.23) in the placebo groups. A mean score of 4.37 (± 0.478) and 2.25 (± 0.736) were observed in the placebo and the vaccinated groups infected with BA.2 respectively. The histopathological scoring is given as supplementary data (Supplementary table 1).

## Discussion

We compared the protective efficacy of COVAXIN® following 2 and 3 dose immunizations against the Delta variant and also studied the efficacy of COVAXIN® against Omicron variants in hamster model. In the Delta infection study, where we compared the protective response between the 2 and 3 dose regimens, we could observe the advantage of the booster dose vaccination in the protection. Although the neutralizing antibody levels were comparable among the groups, lung disease severity was found more reduced after the 3 dose vaccination. The virus shedding and viral organ load were considerably reduced in both the 2 dose and 3 dose immunized animals indicating the vaccine efficacy against Delta variant. The difference in the antibody response could be probably due to the individual animal variation in response as the animals in the 2 dose and 3 dose groups were different. This also points to the role of other components of immune system in protection. We have not assessed the cellular immune response in the present study due to the limitation of availability of reagents specific for the Syrian hamsters. COVAXIN® was found to induce robust immune memory and Th1 skewed response in previous studies.^23,24^ The vaccine was also found to elicit a substantial fraction of T follicular cells which in turn can aid in long term humoral immunity.^23^ The proportion of the antibody secreting memory B cells in the 3 dose COVAXIN® recipients were found more compared to the 2 dose recipients.^10^

For COVAXIN®, an effectiveness of 69% has been reported against severe COVID-19 and 50% against symptomatic COVID-19 for Delta variant in humans after 2 dose vaccinations.^25^ Another vaccine effectiveness study demonstrated 67% effectiveness of the COVAXIN® against severe disease during the Delta variant wave.^25^ The reduced effectiveness of vaccine for prevention of breakthrough infection is also demonstrated.^26^ These studies were performed after a 2 dose vaccine regimen. Decline of neutralizing antibody response was observed after 6 months post 2 dose COVAXIN® immunization, however, persistent AIM+ SARS-CoV-2 specific CD4+ and CD8+ T cell memory phenotype was observed. Third booster dose led to pronounced increase in the neutralizing response against homologous and heterologous SARS-CoV-2 variants in humans as reported in a double-blind, randomised controlled phase 2 clinical trial.^10^ The interval between the 2^nd^ and the 3^rd^ dose vaccination was short in the current study, unlike the real life scenario, where booster doses are recommended after 6 months or after considerable decline in the antibody levels. The booster dose of COVAXIN® was found to improve the neutralizing antibody response against the VoCs including Delta and the Omicron.^27^ Vaccine effectiveness reported for an inactivated COVID-19 vaccine, Coronavac also shows that the three dose immunization give better protection in terms of disease outcomes.^28^ The current data shows that boosting of the immune response tend to improve the vaccine effectiveness for disease severity.

Vaccine efficacy data of 2 dose/3 dose COVAXIN® against Omicron variant in humans is not available. Our findings demonstrate the protective response of the vaccine against the Omicron variant. Limited or no body weight loss and reduced lung disease severity was observed in the vaccinated animals. The disease severity in the COVAXIN® immunized animals were found reduced in spite of this less neutralizing antibody levels following the Omicron variant challenge. The breakthrough cases sera as well as COVAXIN® booster dose vaccinee sera demonstrated better neutralizing titres against Omicron and other VOCs in comparison to the two dose vaccinee sera.^20,27^ Vaccine effectiveness of 50% was reported against symptomatic and severe disease by Omicron variant within 3 months of the first two doses of Coronavac and a booster dose of the vaccine improved the effectiveness.^28^ The vaccine effectiveness studies with the 2 dose regimen of the primary series of mRNA vaccines (Spikevax, Comirnaty), inactivated vaccine (Corona vac), vectored vaccines (Vaxzevria, Ad26.COV2.S.) conducted in 11 countries against Omicron variant has shown reduced effectiveness for disease severity, symptomatic disease, and infection.^12^ Booster vaccination was found to considerably improve the vaccine effectiveness in these studies.^12^ Here we have used 3 dose regimen for Omicron studies considering these observations.

The protection observed might be as a result of the cellular response induced too. In humans, it is demonstrated that the magnitude of immune responses diminishes post 2 dose COVAXIN® over time, whereas booster dose vaccinations enhance the neutralizing antibody responses against homologous and heterologous variants.^10,20^ Apart from B cell mediated immunity, the SARS-CoV-2 specific central and effector memory cells which can aid in cytotoxic function (CD8+ TEMRA phenotype) has been demonstrated in the COVAXIN® recipients.^23^ These properties could have attributed to the rapid reduction of viral load after challenge. Although marked difference was not observed in the histopathological cumulative score of the BA.1.1 infected groups, the mean score of the vaccinated groups were considerably lower compared to our earlier studies on pathogenicity of BA.1.1 in hamster.^29^ There was a limitation of sample size also in the present study.

The evidence from the present in vivo study shows that COVAXIN® booster immunization tends to broaden the protective immune response and reduce disease severity against the Delta and Omicron variant infection.

## Methods

### Study design

The experiments were conducted as per the approval of Institutional Animal Ethics Committee (Approval number: NIV/IAEC/2020/MCL/10) and Institutional Biosafety Committee (Approval number: NIVIBSC/30.05.2020/01) and were performed according to the guidelines of Committee for the Purpose of Control and Supervision of Experiments in Animals (CPCSEA, 2018). Seventy two, 8-10 week old, female Syrian hamsters (*Mesocricetus auratus*) procured from a CPCSEA licensed breeding facility was used in the study. Syrian hamsters were housed in the individually ventilated cages with access to *ad libitum* food and water in the containment facility. The animals were acclimatized to the laboratory conditions for a period of 7 days before the experiments. The animals were observed daily for their activity and a weight loss of 15% was set as humane end point for the study.

### Virus

SARS-CoV-2 variants, Delta (EPI_ISL_2400521), BA.1.1 (EPI_ISL_8542938), BA.2 (EPI_ISL_12667336) and B.1 (EPI_ISL_420546) isolated from a patient’s throat/ nasal swab sample in Vero CCL-81 cells was used for the study.

### COVAXIN® immunization and virus challenge

In the first set of experiments, 40 hamsters were divided into 4 groups of 10 animals each. Animals of two groups (Group 2 and 4) were immunized intramuscularly with 0.2 ml of COVAXIN®. Group 2 received 2 doses of COVAXIN® on day 0 and 14 and group 3 received 3 doses of COVAXIN® on Day 0, 14 and 35. Group I and 3 were given 2 and 3 doses of 0.2 ml of phosphate-buffered saline respectively. The animal caretakers and the technical staff were blinded about the group allocation. Blood samples were collected on day 28 from hamsters which received two doses of COVAXIN® and on day 48 from hamsters which received 3 doses of vaccine and were assessed for IgG and neutralizing antibody response. The hamsters of Group 1 and 2 were challenged with 0.1 ml (10^5^ TCID50/0.1 ml) of SARS-CoV-2 Delta variant on day 35 and the group 3 and 4 were inoculated with Delta variant on day 60. Following the virus challenge the hamsters were monitored for the body weight changes and the viral RNA shed through throat swabs and nasal wash on 1,3,5 and 7 DPI. Five hamsters /group were sacrificed on day 3 and day 7 and lungs and nasal turbinates were collected to understand the viral load. A portion of the lungs sample was immersion fixed in 10% formalin for histopathological investigations.

In the second set of experiments, 36 Syrian hamsters of either sex were used. Sixteen hamsters were immunized with 3 doses of COVAXIN® and the other 16 were given placebo injections of phosphate buffered saline under isoflurane anaesthesia. Eight hamsters from the COVAXIN® immunized group and eight from the placebo group were infected with 0.1 ml of SARS-CoV-2 BA.1.1 variant (1.9 × 10^5^ TCID50/ml) intranasally on day 56 and the rest of the hamsters were infected with BA.2 variant (1.8 × 10^5^ TCID50/ml) on day 70 intranasally. The hamsters were observed for body weight changes post virus infection. Nasal wash and throat swab samples were collected on 1,3,5 and 7 DPI. Four hamsters each were sacrificed on day 3 and 7 post infection and blood, lungs and nasal turbinates were collected to understand the viral load and immune response. Lungs samples were collected for histopathological investigations. Immunization, blood collection and virus inoculation were performed under isoflurane anaesthesia and the euthanasia with isoflurane overdose. The ARRIVE guidelines checklist is provided as a supplementary file.

### Anti-SARS CoV-2 IgG and S1-RBD ELISA

The anti-SARS-CoV-2 IgG and S1-RBD ELISA were performed as described earlier.^30^

### Plaque Reduction Neutralization test

The assay was performed using B.1, Delta, BA.1.1 and BA.2 variants as described earlier.^31^

### Quantitative Real-time RT-PCR

Nasal wash and throat swab samples collected in 1ml viral transport medium and weighed organ samples (lungs, nasal turbinate) homogenized in 1 ml sterile tissue culture media were used for RNA extraction. MagMAX™ Viral/Pathogen Nucleic Acid Isolation Kit was used for the procedure as per the manufacturer’s instructions. Published primers targeting E gene was used for viral RNA estimation and tragetting N gene were used for the subgenomic RNA estimation.^32,33^

### Virus titration in Vero cells

The lungs, nasal turbinate and nasal wash samples were used for virus titration in Vero CCL81 cells. Twenty four**-**well plate with Vero CCL81 monolayers were incubated for 1 hour after addition of the hundred microlitre of the sample. The plate was washed after removal of the media and was incubated again with maintenance media containing 2% FBS in a CO2 incubator. The plates were examined for any cytopathic effects daily.

### Histopathology

The formalin fixed lung tissue samples were processed by routine histopathological techniques for hematoxylin and eosin staining.^34^ The lung lesions were scored for vascular lesions like congestion and hemorrhages, perivascular inflammatory cell infiltration, bronchiolar changes like epithelial loss/necrosis, exudates, inflammation, alveolar changes like hyperplasia, hyaline formation, edema, emphysema and inflammatory cell infiltration. The sections were scored on a scale of 0 to 4 and were averaged and plotted.

### Data analysis

For analysis of the data, Graphpad Prism version 9.4.3 software was used. Mann-Whitney test was performed between the placebo and the vaccinated groups. The p-values less than 0.05 were considered to be statistically significant.

## Data availability

All the data related to the experiments are available in the manuscript and supplementary data.

## Author Contributions

PDY and SM conceived and designed the study. KM and BG prepared and provided the vaccine formulations. SM performed the animal experiments. PDY, SM, ASA, GS and GD performed the laboratory work planning. GD and GS performed the neutralization assays. ASA and RJ performed the ELISA. AK and KW assisted in animal experimentation, data collection and performed sample collection. HD performed the virus neutralization assay. AK, KW, JY and PG performed the sample processing in the laboratory. PDY and SM have drafted the manuscript. KM, BG and PA substantively revised the manuscript. All authors reviewed the manuscript and agree to its contents.

