## Supplemental data for "Protective efficacy of COVAXIN® against Delta and Omicron variants in hamster model"

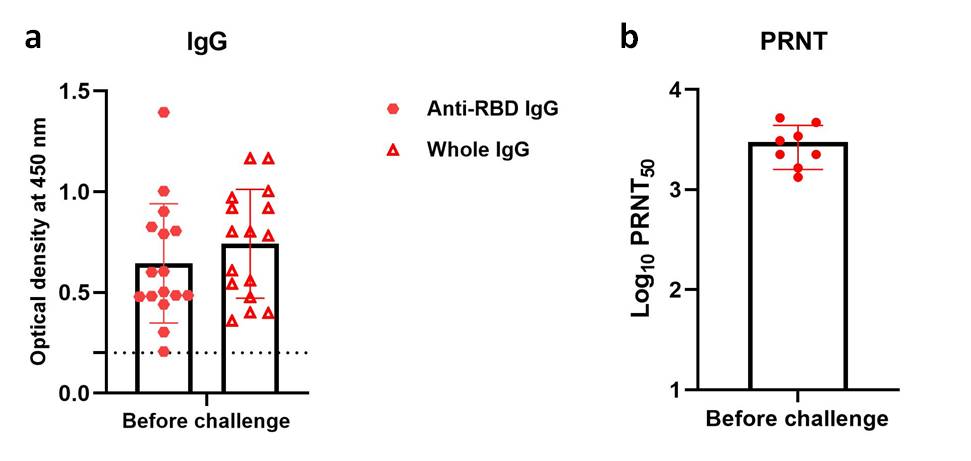

**Supplementary figure 1: Immune response in 3 dose immunized animals of the Omicron group before challenge.** a)Anti-SARS-CoV-2 IgG against whole virion and anti-SARS-COV-2 RBD IgG in hamsters before Omicron virus challenge (n=16 /group). The bars represent mean and standard deviation. b) Neutralizing antibody titres of the hamster sera against B.1 variant before virus challenge. n=10, bars represent geometric mean and standard deviation.

**Supplementary table 1:** Histopathological scoring of the lung lesions of hamsters on 7 days post infection.

| **Group** | **Animal number** | **Vascular lesions**  **(Congestion, haemorrhages)** | **Bronchial lesions**  **(loss of epithelium, exudation, degeneration)** | **Alveolar lesions (Consolidation, septal thickening, hyperplasia)** | **Edematous and emphysematous changes** | **Inflammatory infiltration in alveolar interstitium** | **Peribronchial and perivascular infiltration** |
| --- | --- | --- | --- | --- | --- | --- | --- |
| Placebo  (2 dose) + Delta | 1 | +2 | +2 | +2 | +2 | +2 | +2 |
|  | 2 | +2 | +2 | +3 | +2 | +2 | +3 |
|  | 3 | +3 | +2 | +2 | +1 | +2 | +2 |
|  | 4 | +2 | +2 | +3 | +2 | +3 | +3 |
|  | 5 | +2 | +2 | +2 | +1 | +2 | +2 |
| Covaxin  (2 dose) + Delta | 1 | +2 | +1 | +3 | +1 | +2 | +2 |
|  | 2 | +1 | +1 | +2 | +1 | +2 | +2 |
|  | 3 | +1 | 0 | +2 | 0 | +1 | +2 |
|  | 4 | +1 | +1 | +1 | 0 | +1 | +2 |
|  | 5 | +3 | +1 | +3 | +1 | +1 | +2 |
| Placebo  (3 dose) + Delta | 1 | +3 | +3 | +4 | +2 | +3 | +3 |
|  | 2 | +2 | +2 | +3 | +2 | +3 | +3 |
|  | 3 | +3 | +2 | +4 | +2 | +3 | +3 |
|  | 4 | +4 | +2 | +4 | +2 | +3 | +3 |
|  | 5 | +2 | +2 | +3 | +1 | +3 | +3 |
| Covaxin  (3 dose) + Delta | 1 | +1 | +1 | +1 | 0 | +1 | +1 |
|  | 2 | +1 | +1 | +1 | 0 | +1 | +1 |
|  | 3 | +1 | +1 | +1 | 0 | 0 | +1 |
|  | 4 | +1 | +1 | +1 | 0 | 0 | +1 |
|  | 5 | +1 | +1 | +1 | 0 | +1 | +1 |
| Placebo  (3 dose) + BA.1.1. | 1 | +1 | 0 | +2 | 0 | +2 | +2 |
|  | 2 | +2 | +1 | +2 | +2 | +2 | +3 |
|  | 3 | +3 | +3 | +4 | +2 | +3 | +3 |
|  | 4 | +2 | +2 | +2 | +2 | +2 | +3 |
| Covaxin  (3 dose) + BA.1.1. | 1 | +1 | 0 | +2 | 0 | +2 | +2 |
|  | 2 | +2 | +1 | +2 | +2 | +2 | +2 |
|  | 3 | +2 | +2 | +2 | +1 | +2 | +2 |
|  | 4 | +2 | +1 | +2 | +1 | +2 | +2 |
| Placebo  (3 dose) + BA.2 | 1 | +3 | +2 | +3 | +1 | +3 | +3 |
|  | 2 | +3 | +3 | +4 | +2 | +3 | +3 |
|  | 3 | +3 | +3 | +4 | +2 | +3 | +3 |
|  | 4 | +3 | +3 | +3 | +2 | +3 | +3 |
| Covaxin  (3 dose) + BA.2 | 1 | +2 | +2 | +2 | +1 | +2 | +2 |
|  | 2 | +2 | +2 | +2 | +2 | +2 | +2 |
|  | 3 | +1 | 0 | +1 | 0 | +1 | +2 |
|  | 4 | +2 | 0 | +2 | +1 | +2 | +2 |
